# Palytoxin Evokes Reversible Spreading Depolarization in the Locust CNS

**DOI:** 10.1101/2024.06.11.598415

**Authors:** Yuyang Wang, Rachel A. Van Dusen, Catherine McGuire, R. David Andrew, R. Meldrum Robertson

## Abstract

Spreading depolarization (SD) describes the near-complete depolarization of CNS neural cells as a consequence of chemical, electrical, and metabolic perturbations. It is well-established as the central mechanism underlying insect coma and various mammalian neurological dysfunctions. Despite significant progress in our understanding, the question remains: which cation channel, if any, generates SD in the CNS?

Previously, we speculated that the sodium-potassium ATPase (NKA) might function as a large-conductance ion channel to initiate SD in insects, potentially mediated by a palytoxin (PLTX)-like endogenous activator. In the current study, we evaluate the effectiveness and properties of PLTX as an SD initiator in *L. migratoria*. Whereas bath-applied PLTX failed to ignite SD, direct injection into the neuropil triggered SD in 57% of the preparations. Notably, PLTX-induced SD onset was significantly more rapid compared to ouabain injection and azide controls, though their electrophysiological features remained similar. Furthermore, PLTX-induced SD was recoverable and resulted in a greater frequency of repetitive SD events compared to ouabain.

Surprisingly, prior PLTX treatment disrupted the onset and recovery of subsequent SD evoked by other means. PLTX injection could attenuate the amplitude and hasten the onset time of azide-induced SD. Such an effect is associated with a complete inhibition at higher doses of subsequent anoxic SD induced through azide treatment or submersion.

These results show that PLTX can trigger repetitive and reversible SD-like events in locusts and simultaneously interfere with anoxic SD occurrence. We suggest that the well-documented NKA pump conversion into an open non-selective cationic channel is a plausible mechanism of SD activation in the locust CNS, warranting additional investigations.

## Introduction

Spreading depolarization (SD) describes the propagation of repetitive waves of neural cell depolarization in central nervous systems (CNS). SD is characterized by an almost complete loss of membrane potential, large changes in intra- and extracellular ion concentrations, and the silencing of electrical activities (Rossini and Bigiani, 2011: Andrew, Hartings, et al., 2022). The prevalence of SD has been extensively documented in vertebrates and invertebrates and may be an emergent property of a high-performance CNS protected by a diffusion barrier (Robertson et al., 2020). Despite important differences in neuroarchitecture, the similarities in trajectory and triggers suggests that insect and mammalian SD may share a common underlying process (Shuttleworth et al., 2020; Spong et al., 2016).

Although extracellular potassium ([K^+^]_o_) and excitatory neurotransmitters promote SD activation and propagation (Somjen, 2001), there is consensus that neither directly initiate SD and the true biological activator remains unknown (Andrew, Hartings, et al., 2022). Nevertheless, increased neuronal activity or impaired clearance elevates [K^+^]_o_, shifting the K^+^ reversal potential and depolarizing neurons via voltage-gated ion channels. Concurrently, excitatory neurotransmitter build-up opens ligand-gated ion channels. Both of these effects increase tissue susceptibility, but an additional mechanism is required to initiate the all-or-none event of SD. Once initiated, sodium flux (but not through voltage-gated Na^+^ channels) has been suggested as the primary driver of cellular depolarization (Herreras & Makarova, 2020). Irrespective of the relative contributions of ions, the membrane conductance must increase initially to permit ion currents to kickstart the positive feedback process in SD generation. Such a conductance increase is likely mediated by an unknown voltage-independent, non-specific Na^+^/K^+^ channel (Czéh et al., 1993; Somjen et al., 2009).

The Zoanthid palytoxin (PLTX) is one of the world’s most potent poisons and is documented as disrupting sodium/potassium ATPase (NKA) gating (Artigas & Gadsby, 2003, 2004; Rossini & Bigiani, 2011), thereby increasing ion conductance in mammalian cardiac myocytes, squid giant axons, and cockroach interganglionic connectives (Kockskämper et al., 2004; Pichon, 1982). Thus a plausible candidate for the SD initiating channel is the NKA, which has been proposed to convert to a cation channel under ischemic conditions to drive SD (Andrew et al., 2013; Andrew, Hartings, et al., 2022). An endogenous SD activator molecule (SDa) could mimic the effect of PLTX that drives SD generation (Andrew et al., 2013). Compelling evidence for the existence of such an SDa is that perfusate from brain slices that had undergone SD could trigger SD in naïve slices, independent of K^+^, pH, and glutamate levels (Hellas, 2023; Lee, 2020). The effect of a putative SDa can be enhanced by 0.01 nM PLTX priming of tissue slices (Lowry et al., 2021). Further, higher nanomolar concentrations of PLTX induce SD-like events similar to oxygen-glucose deprivation (OGD) in rat brain slices (Brisson et al., 2014; Gagolewicz et al., 2016), where patch recordings also show the activation of a novel channel during OGD with properties similar to those evoked by PLTX (Gagolewicz & Andrew, 2017). The pump-to-channel conversion also establishes a direct link between pump arrest and SD initiation, especially in well perfused nervous tissue with ample ATP supply.

In this study, we investigated PLTX’s ability to induce SD in the locust CNS *in vivo* via bath application into the thoracic cavity, and microinjection into the ganglia. We identified SD as abrupt negative DC shifts of the extracellular potential in the neuropil by recording the transperineurial potential (TPP) across the hemolymph-brain barrier (HBB). We also characterized the impact of PLTX on neural function and TPP trajectories. We then compared these effects of PLTX to traditional SD triggers such as ouabain and azide.

## Methods

### Animals

Gregarious *Locusta migratoria* were reared in a crowded colony in the Animal Care facility of Biosciences Complex at Queen’s University, Kingston. A 12-hour light-dark cycle was maintained, during which the ambient temperature was 30 °C under light and room temperature when dark. Animals were fed each day with a diet of fresh wheatgrass, granular yeast, milk protein, and bran. Locusts 3-6 weeks post-imaginal moult were selected for experiments, and randomly assigned to treatments groups. Animals were transported in well-ventilated containers between the colony and lab spaces. Roughly equal numbers of males and females were used in palytoxin (PLTX) bath experiments. Animal sex for injection experiments was as follow, control: 4 males and 10 females; ouabain (OUA): 6 males and 9 females; PLTX:11 males and 35 females. N2 anoxia experiments consisted of 5 males and 4 females. More females were used in injection experiments, as smaller size of male ganglia resulted in interference with injector and DC electrode placements. Data from both sexes were pooled for reporting.

### Pharmacology

Three final concentrations of PLTX were used for bath experiments: 100 nM (4 preparations), 500 nM (8 preparations), and 1000 nM (4 preparations). Note that these concentrations are 10-100 fold higher than used in rodent brain slices (Andrew et al., 2013). PLTX solution was prepared from a single 100 µg palytoxin vial with ddH_2_O, and made up into 10 uM stocks (Wako Chemicals USA, Richmond, VA). For bath application during electrophysiology, frozen aliquots of 10 μM PLTX stock were diluted to final concentrations in standard locust saline (147 mM NaCl, 10 mM KCl, 4 mM CaCl_2_, 3 mM NaOH & 10 mM HEPES buffer (pH = 7.2)) before each experiment. 10 mM sodium azide in standard locust saline was used for control and azide bath following PLTX injection. The injection experiments used 10 μM PLTX or 10 mM ouabain (OUA, Ouabain octahydrate, Sigma-Aldrich, St. Louis, MO, USA) in standard saline.

### Semi-Intact Dissection

See **Fig.1A** for schematic of semi-intact preparation. The legs and the dorsal pronotum were removed. An incision along the dorsal midline was made from the seventh abdominal segment to the head capsule. The thoracic cavity was then exposed by pinning the animal to a cork board.

**Figure 1.**
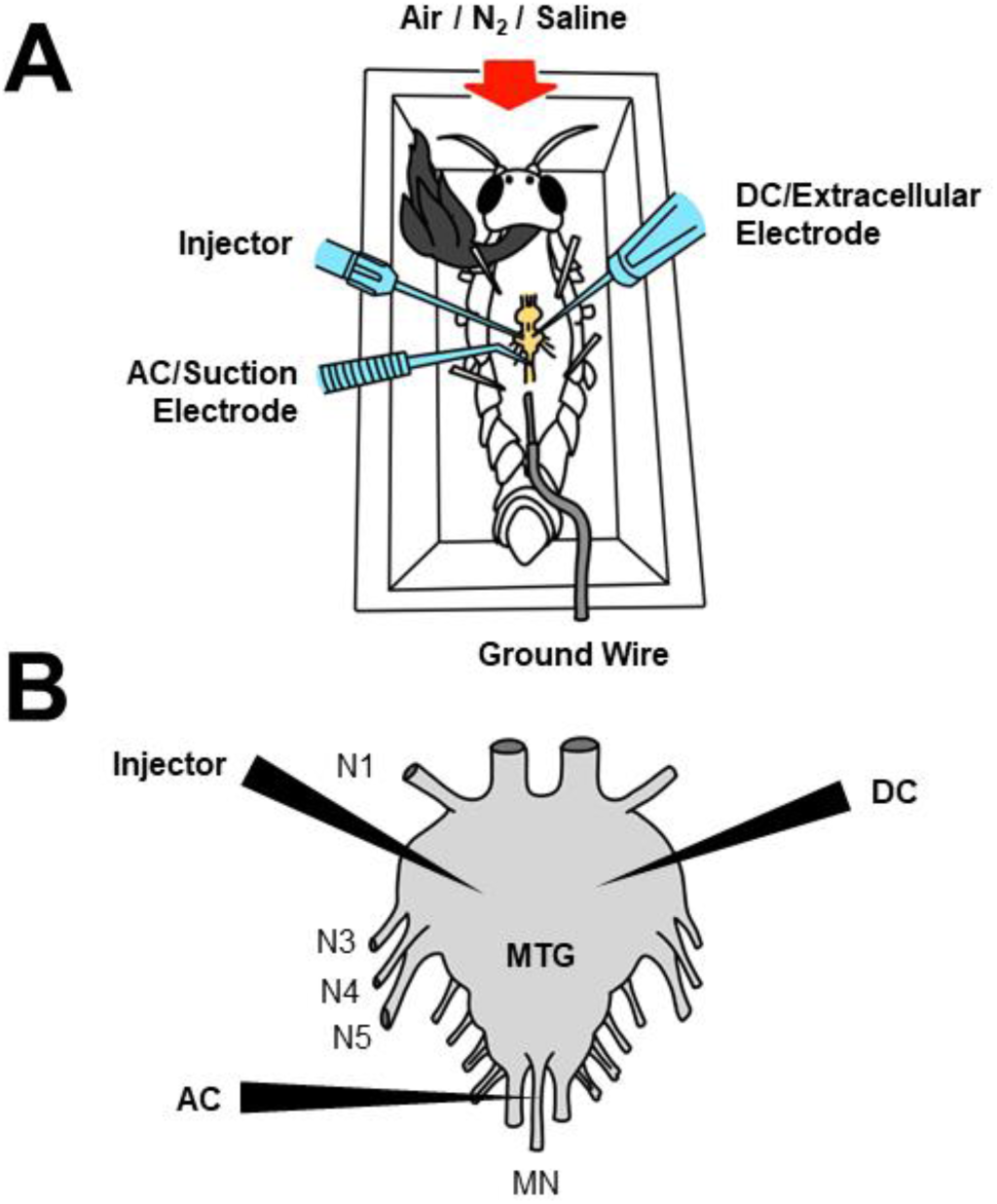
Experimental setup. **A**: schematic of semi-intact animal preparation in a Plexiglas chamber used for submersion and N_2_ anoxia. **B**: enlarged dorsal view of the metathoracic ganglion showing electrode and injector placement. DC electrode and injector penetrate into the neuropil, whereas AC electrode attaches to the median nerve (**MN**). Peripheral nerves (**N3, 4, 5**) on both sides were cut near the roots in PLTX bath experiments to improve drug access. Refer to Methods for details.

The air sacs, fat bodies, and gonads were then removed. The gut was severed caudally and pulled to one side. The ventral diaphragm was subsequently removed to allow access to the metathoracic ganglion (MTG) and associated nerve roots. Finally, the ventral longitudinal tracheae surrounding the MTG were clipped and removed, while leaving the ventral air sacs intact. Standard saline was dripped into the thoracic cavity during dissection to prevent desiccation. The nerve roots N3, 4, and 5 were cut near the ganglion surface bilaterally in 4 PLTX bath preparations, and the MTG was mechanically desheathed in 2 preparations, to improve PLTX access to the neuropil.

### Extracellular Recording Setup

The placement of electrodes and injector is shown in **Fig. 1B**. The semi-intact animal was grounded with a chlorided silver wire in contact with the bathing saline. An extracellular DC electrode pulled from 1 mm filamented glass capillary (World Precision Instruments, Sarasota, FL, USA) to a tip resistance of 5 MΩ was used to measure transperineurial potential (TPP). The electrode and holder were backfilled with 500 mM and 3 M KCl, respectively. The electrode tip penetrated the perineurium and sub-perineurium to access the neuropil. Signals were amplified with a DC amplifier (Model 1600, A-M Systems, Sequim, WA, USA). Calibration was performed before and after each experiment in grounded bathing saline to account for baseline drift. An AC suction electrode was used to measure nerve activity and ventilatory action potentials on the dorsal median nerve of MTG. The AC electrode was pulled using 1 mm non-filamented glass capillary to a tip resistance of 5 MΩ; the tip was then carefully broken with forceps to match the diameter of the median nerve. A syringe was used to fill the AC electrode with bathing saline and form a suction seal around the nerve surface. The signals were passed to a differential AC amplifier (model 1700, A-M Systems, Sequim, WA, USA). The analog signals from both amplifiers were digitized (DigiData 1322A, Molecular Devices, San Jose, CA, USA) and recorded using Axoscope 10.7 (Molecular Devices, San Jose, CA, USA).

### Recording of SD

SD occurrence was derived from the trajectory of transperineurial potential (TPP), which is the extracellular DC potential measured from the neuropil. TPP represents the difference between the adglial (neuropil facing) and basolateral (hemolymph facing) membrane potentials of the barrier-forming perineurial glial cells (Schofield & Treherne, 1984). Sustained SD refers to an initial abrupt negative DC shift without recovery. The term “Terminal SD” was not used because we did not confirm whether SD could be reversed by washing the preparation, which allows recovery for azide treatment (Rodgers et al., 2010). Repetitive SD refers to periodical SDs that recover and spontaneously re-ignite. Recoverable SD refers to a single SD that recovers, without downstream repetitive events within the time course of the experiment.

In PLTX bath experiments, animals were dissected in a Plexiglas chamber (5 x 2.5 x 2 cm). Following dissection and electrode placement, the bathing standard saline in the thoracic cavity was replaced with PLTX saline. The experiment then proceeded for 1 hour, at the end of which the animal was fully submerged with saline for 10 minutes to induce anoxic SD. A 5-minute reoxygenation period followed, where saline was drained and the animal was allowed to ventilate in air to recover.

In injection experiments, animals were dissected on an open cork board to allow injector access. A Nanoject III microinjector (Drummond Scientific, Broomall, PA, USA) was used to deliver 23 nL boluses into the ganglion on a slow speed setting (92 nL/s). The injection pipette was pulled from manufacturer supplied 1.14 mm glass capillaries to a tip resistance of approximately 5 MΩ. The tip was then carefully broken off with forceps, and the pipette backfilled with either standard saline, OUA, or PLTX. The DC electrode and injection pipette were positioned before the start of each experiment; an AC suction electrode was not used due to equipment space limitations.

Control animals received 46-115 nL of standard saline injection to rule out SD induction through mechanical disturbance; this was then followed by an azide bath to induce SD. Ouabain animals received 115 nL injections to induce SD. Palytoxin animals received incremental boluses, starting from 23 nL until SD was induced or the 45 minutes experiment endpoint. The PLTX animals were subjected to an azide bath at the end of the experiments to induce anoxic SD, except for those that experienced sustained SD from PLTX injection.

In N_2_ anoxia experiments, animals were dissected in a similar Plexiglas chamber as bath experiments, which was connected to a N_2_ tank and an air pump for ventilation. The chamber was sealed with cellophane tape following electrode placement. N_2_ anoxia was introduced; 10 minutes after the abrupt negative DC shift, the N_2_ was switched off and the air pump reoxygenated the chamber. After the first anoxia-reoxygenation cycle, the bathing standard saline was replaced by 500 nM PLTX saline. The animals were then subjected to repeated N_2_ anoxia with 10 minutes interval.

The abrupt negative DC shift signalled an SD event. TPP positivity is the peak amplitude of positive DC shift after PLTX treatment. SD latency refers to the times taken from the start of treatment to the mid-amplitude point of the negative DC shift. SD amplitude represents the voltage difference between steady state TPP before SD and the minimum point of TPP during SD. Onset slope is measured as the tangential linear slope of TPP descent at the mid amplitude point of SD. In bath and injection experiments, SD induced within 30 minutes of treatment start was counted. Only the first SD of a repetitive event was used for analysis.

### Data Analyses

Sigmaplot 13 was used to perform statistical analysis and generate figures. Data normality and equal variance were assessed by Shapiro-Wilk and Brown-Forsythe tests, respectively. For normally distributed data, student’s t-test (paired and unpaired) and One-way ANOVA were used to compare group means. Otherwise, Mann-Whitney ranked sum tests and One-way ANOVA on ranks were used to compare medians. Dunn’s method was used as a post hoc pairwise analysis to determine significance between groups for non-parametric data. Alpha was set at p = 0.05. All data were presented as boxplots showing medians and IQR, with whiskers to the 5th and 95th percentile. Samples closest to the group mean were chosen as representative traces presented in figures. The recordings were processed using Clampfit 10.7 (Molecular Devices, San Jose, CA, USA) to convert images and removed abrupt electrical artifacts (<1 ms).

## Results

### PLTX Bath Does Not Generate SD, but Modifies Nerve Excitability

To examine the influence of PLTX on CNS functioning, we recorded efferent nerve activity from the metathoracic ganglion (MTG; **Fig. 2Ai**). PLTX had no immediate effect on the frequency of ventilatory motor patterning, which remained unchanged after 5 minutes (Paired t-test, Pre-bath versus Post-bath, n = 8, p = 0.27). Nerve excitability is not immediately eradicated in the presence of PLTX but gradually declined in amplitude (**Fig. 2Aii, Aiii**).

**Figure 2.**
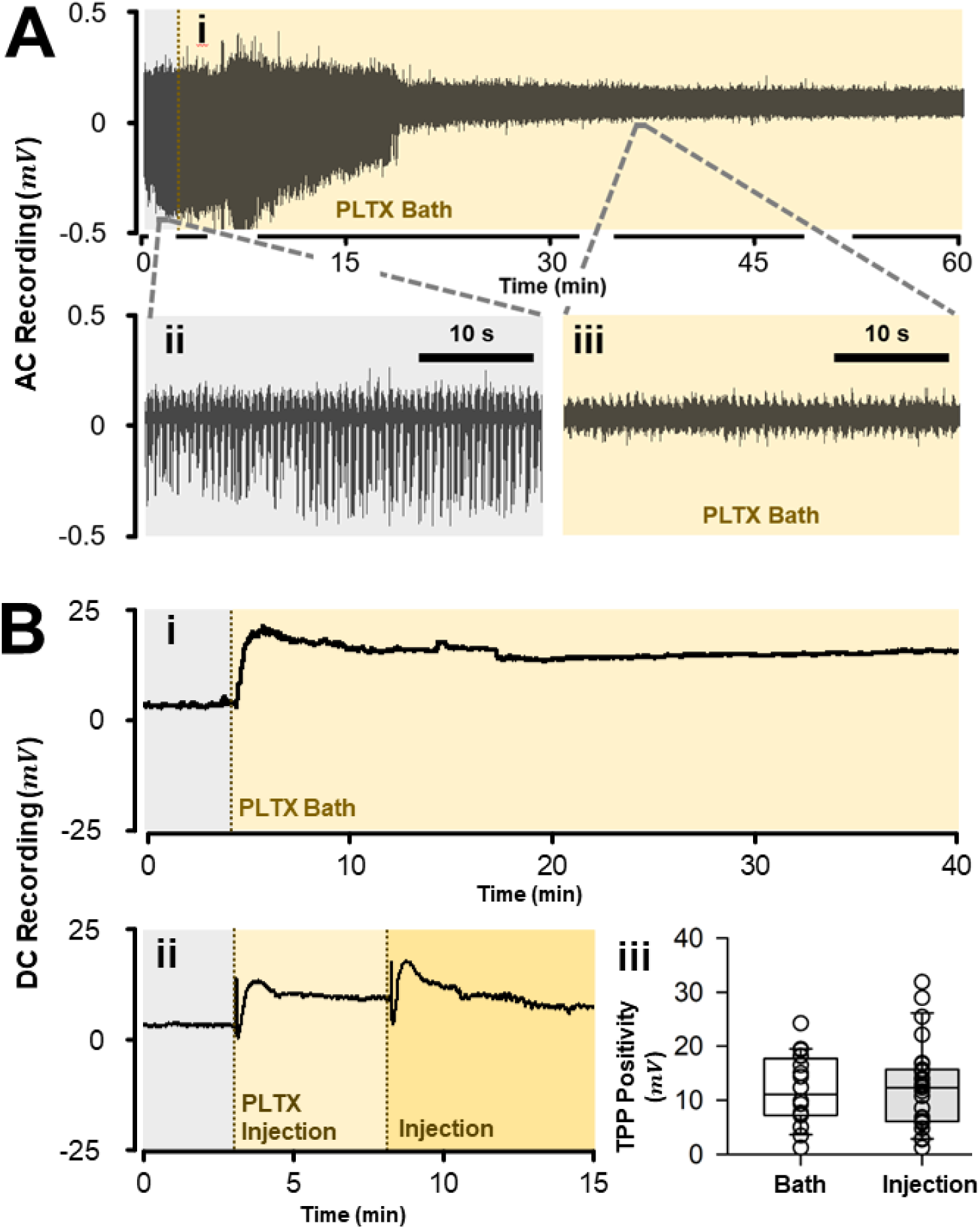
Effects of bath-applied or direct injection PLTX on nerve activity and TPP. **Ai**: representative recording of nerve activity during PLTX bath experiments. **Aii, iii:** the amplitude of the ventilatory action potentials gradually decreases, but some motor patterning persists after 30 mins of PLTX. **Bi:** PLTX bath triggers an immediate and persistent positive shift of TPP. **Bii:** direct PLTX injections into the ganglion generate a similar positive shift. **Biii**: the amplitudes of TPP positivity induced by PLTX is consistent across bath and injection experiments. Grey and yellow backgrounds represent untreated and PLTX-treated time intervals, respectively, with vertical dashed lines indicating the beginning of treatment. Box plots indicate medians and inter-quartile range with whiskers to 5^th^ and 95^th^ percentiles; open circles are individual data points. Refer to the results section for Sample sizes and p-values.

To investigate whether PLTX is an effective SD activator, we subjected semi-intact preparations with locust saline containing 100 to 1000 nM PLTX. Out of 16 preparations, only a single SD-like event was recorded 43 mins after bath application of 500 nM PLTX saline, beyond the 30 mins cut-off (**Table 1**). In contrast to the lack of effect on motor patterning, PLTX immediately generated a sustained positive shift of trans-perineurial potential (TPP), after which the TPP remains stable (**Fig. 2Bi**). Such positivity is also observed in PLTX injection experiments (see next section), and the amplitude appeared similar to bath application (**Fig. 2Bii, Biii**; unpaired Student’s T-test, n = 47, p = 0.792). The TPP positivity presumably represents depolarization of the basolateral membrane of the hemolymph-brain barrier (HBB). It is unknown whether longer incubation times would increase the likelihood of SD onset.

**Table 1.**
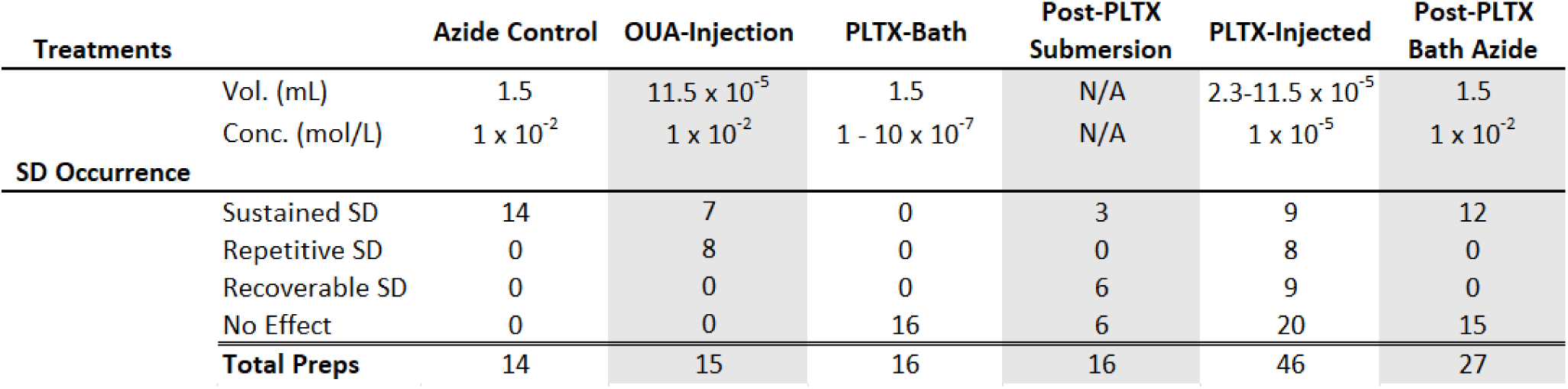
Summary of experiments.

### PLTX Injection into CNS Can Rapidly Induce SD

To bypass the hemolymph-brain barrier and improve neuropil access, we directly injected 10 μM PLTX into the extracellular space of the MTG in 23 nL boluses. Preliminary injection experiments with lower PLTX concentration (0.5-1 μM) could not generate SD. The injection experiments involved 2 sets of controls. First, standard saline was injected in control animals to rule out SD induction through mechanical disturbance from the injector. These control animals were then subjected to azide bath to induce anoxic SD to establish reference SD parameters. Second, another group of animals received direct ouabain (OUA) injection into the ganglia, to control for route of delivery.

57% of the PLTX-injected ganglia generated SD (**Table 1**), whereas 10 mM azide-bath and 23 nL of 10 mM OUA injection controls triggered SD in all preparations. The descriptions of SD generated through various means are documented in **Table 1**, and representative SD waveforms shown in **Figure 3**. Azide consistently triggered an SD that was thereafter sustained throughout the treatment (**Fig. 3Ai**). OUA injection resulted in both sustained and repetitive SD (**Fig. 3Aii, Aiii**). PLTX-induced SD manifested as sustained, repetitive, and single recoverable types (**Fig. 3Aiv, Av, Avi**). The occurrence of different SD events in OUA and PLTX experiments are independent of dosage, as single bolus delivery could trigger all types. On the other hand, 43% of the PLTX injected animals did not show signs of SD-like events (**Table 1**). These preparations are similar to the PLTX bath experiments in that the TPP was stable and maintained above 0 mV over the course of the treatment.

**Figure 3.**
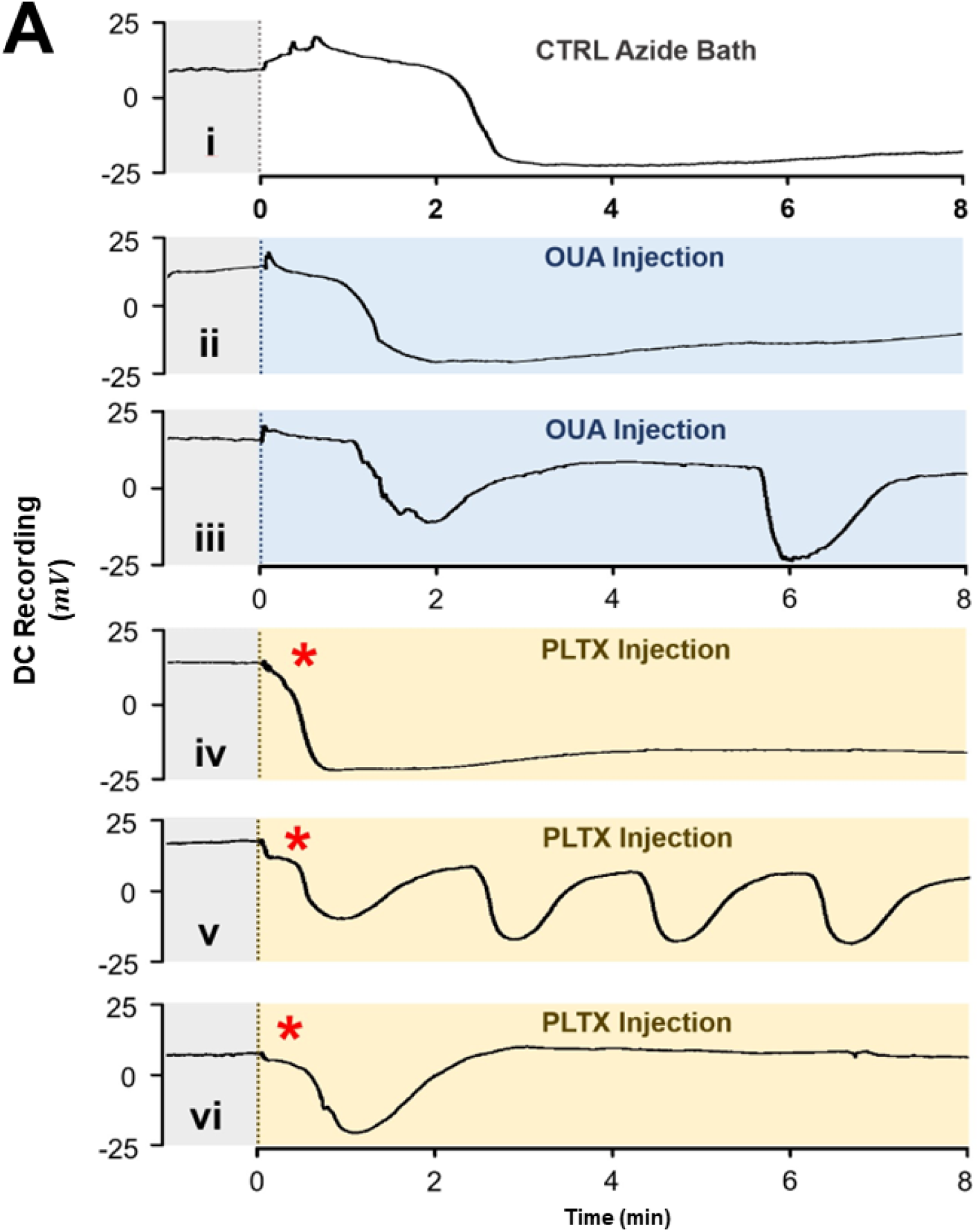
SD waveforms induced by azide, ouabain, and palytoxin. **Ai**: representative recording of azide-induced sustained SD. **Aii, iii**: OUA-induced sustained and repetitive SD. **Aiv, v, vi**: PLTX-induced sustained, repetitive, and single-recoverable SD, respectively. Note the rapid onset of PLTX-induced SD compared to azide and OUA, marked by red asterisks (*). Grey, white, blue, and yellow backgrounds represent untreated, azide, ouabain, and PLTX-treated time intervals, respectively. Vertical dashed lines indicate the beginning of treatment. **CTRL**: azide; **OUA**: ouabain; **PLTX**: palytoxin.

Interestingly, when it occurred, PLTX injection SD had a significantly shorter latency and less variability compared to azide and OUA (**Fig. 4Ai, Bi**; One way ANOVA on ranks, n = 55, p < 0.001; Dunn’s pairwise comparison, OUA versus azide control, p = 0.827; PLTX versus OUA, p < 0.001; PLTX versus azide control, p < 0.001; IQR, Azide = 0.96, OUA = 1.08, PLTX = 0.42), where the negative shift of TPP occurred almost immediately upon delivery (**Fig. 4Ai**).

**Figure 4.**
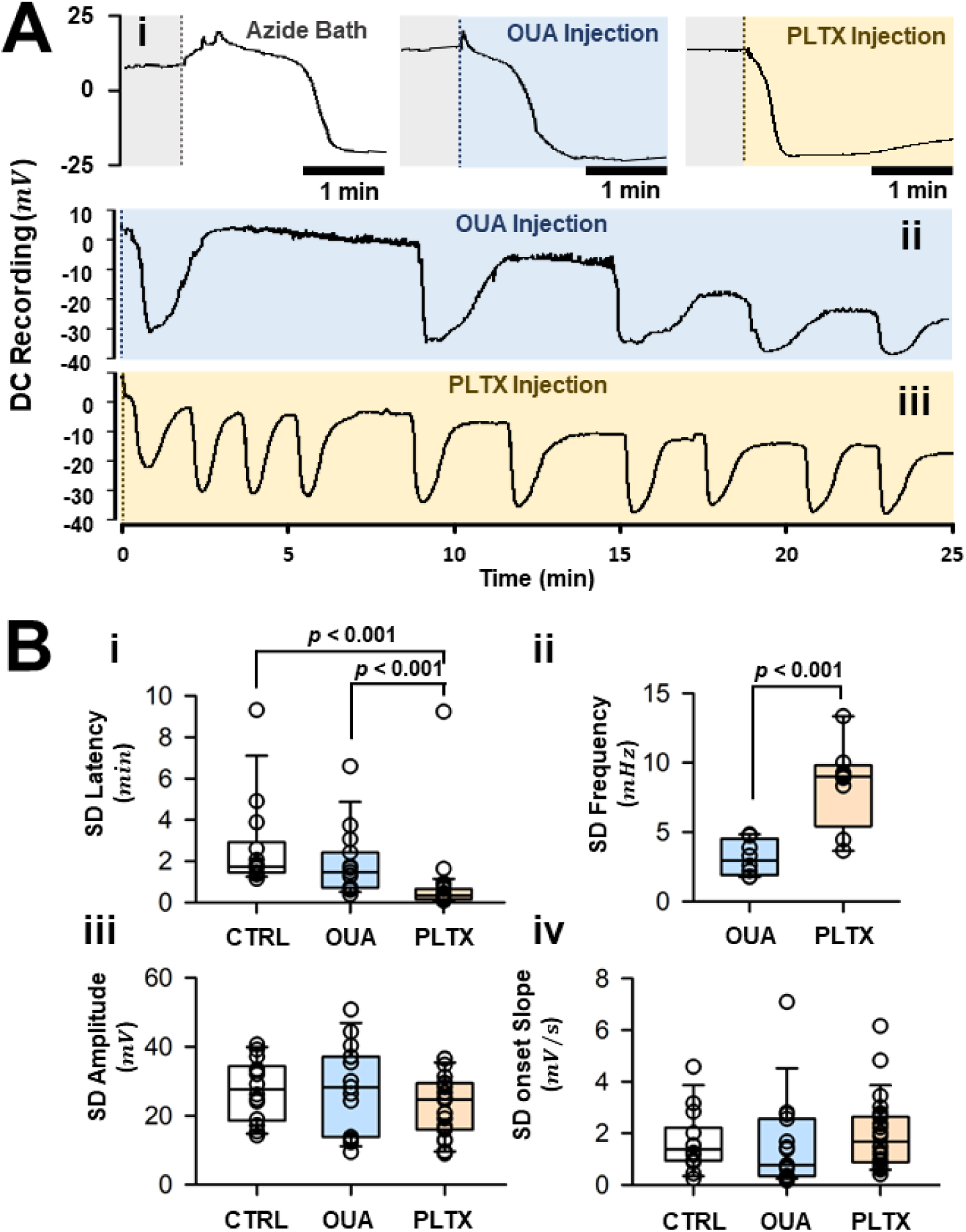
Parameters of PLTX injection SD. **Ai:** representative traces of sustained SD induced via azide bath (left), ouabain injection (middle), and PLTX injection (right). Note the rapid SD onset in the PLTX group. **Aii, iii:** repetitive SD induced by injected PLTX occurs at greater frequency compared to injected ouabain. **Bi, ii**: quantification of SD onset latency and repetition frequency as illustrated in the previous panel. **Biii, iv:** the SD amplitude and onset slope are otherwise similar between azide control, ouabain, and PLTX. Grey, white, blue, and yellow backgrounds represent untreated, azide, ouabain, and PLTX-treated time intervals, respectively. Vertical dashed lines indicate the beginning of treatment. **CTRL**: azide; **OUA**: ouabain; **PLTX**: palytoxin. Box plots indicate medians and inter-quartile range with whiskers to 5^th^ and 95^th^ percentiles; open circles are individual data points. Refer to the results section for Sample sizes and p-values.

Moreover, the mean frequency of repetitive SD was 2.7-fold higher in PLTX groups compared to OUA control (**Fig. 4Aii, Aiii, Bii**; 2 sample Student’s T-test, n = 16, p < 0.001). On the other hand, PLTX injection SD appears similar to azide and OUA control both in amplitude and SD onset slope (**Fig. 4Biii, Biv**; SD amplitude, One way ANOVA, n = 55, p = 0.132; SD onset slope, One way ANOVA on ranks, n = 55, p = 0.346). These findings indicate that, when effective, PLTX can trigger SD in the locust CNS that is rapidly induced and has a higher frequency of repetitive events.

### PLTX Attenuates and Can Eradicate SD

As injected PLTX failed to induce SD in nearly half of the animals, we tested whether these preparations were capable of generating SD by other means. At the end of each bath or injection PLTX experiments, the animals were subjected to submersion or 10 mM azide bath, respectively (**Fig. 5**).

**Figure 5.**
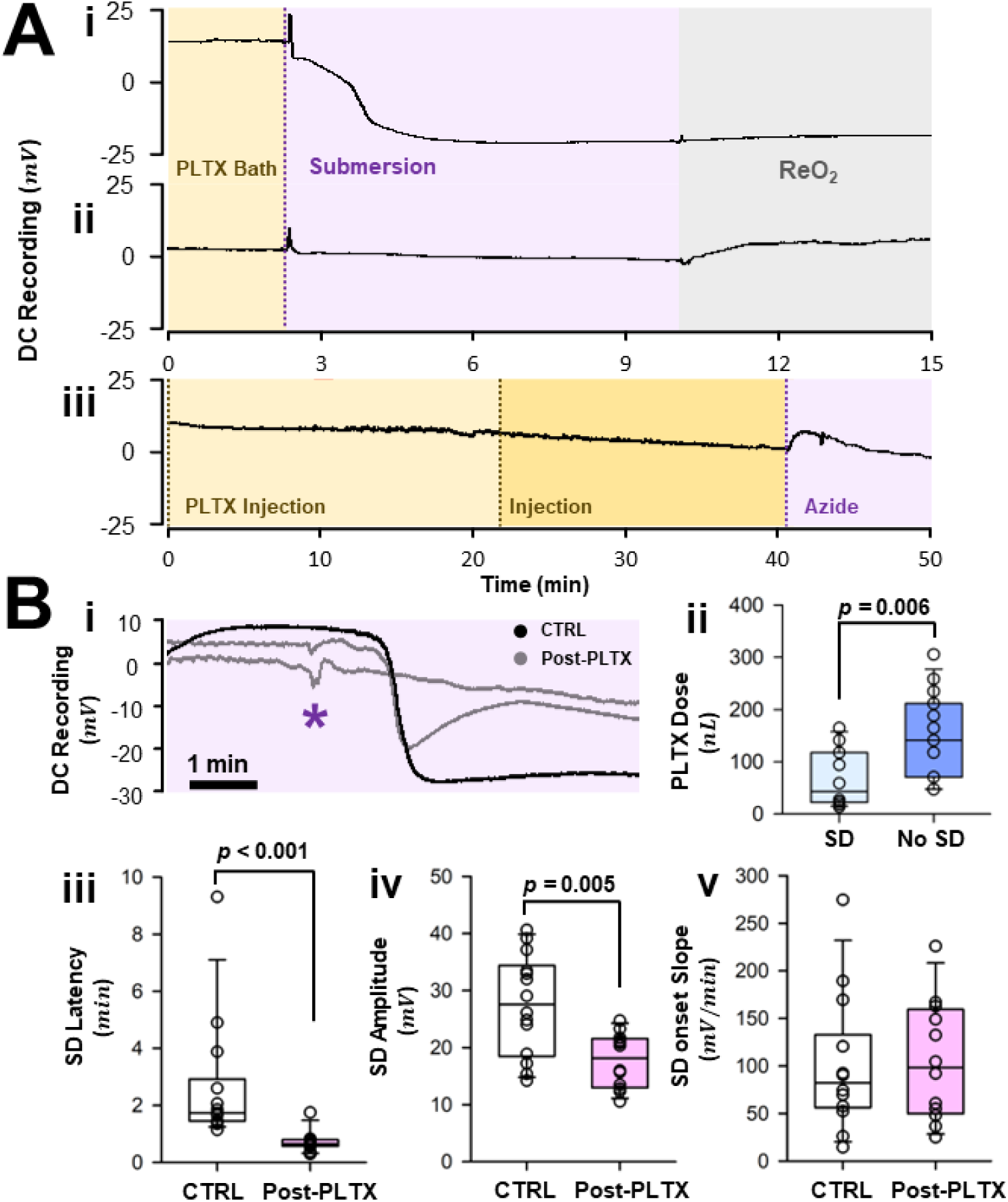
Effects of PLTX on subsequent anoxic SD. **Ai, ii**: prior bath-applied PLTX could affect whole-prep submersion anoxia by either abolishing TPP recovery during reoxygenation, or preventing SD onset. **Aiii**: PLTX injections could similarly prevent subsequent azide-induced SD. **Bi:** overlay of azide-induced SD traces illustrating attenuated and lost negative DC shift in PLTX injection animals. Azide-SD following PLTX treatment also exhibits faster onset. Azide bath for **CTRL** and **Post-PLTX** starts at the beginning of recording and at asterisk (**\***), respectively. **Bii**: the eradication of azide-SD is associated with a higher PLTX injection dosage. **Biii, iv**: quantification of lower SD onset latency and attenuated amplitudes in the PLTX group as illustrated in **Bi**. **Bv**: the SD onset slope is similar between control and PLTX groups. Magenta and yellow backgrounds represent azide and PLTX-treated time intervals, respectively. Grey background indicates reoxygenation. Vertical dashed lines mark the beginning of treatment. **CTRL**: azide; **Post-PLTX**: azide after palytoxin; **ReO_2_**:reoxygenation. Box plots indicate medians and inter-quartile range with whiskers to 5^th^ and 95^th^ percentiles; open circles are individual data points. Refer to the results section for Sample sizes and p-values.

Surprisingly, PLTX could completely inhibit anoxic SD onset. Despite differing success in SD induction, both bath and injected PLTX blocked anoxic SD in a considerable portion of the preparations (**Table 1**). In these animals, the usual abrupt negative DC shift under anoxia was absent (**Fig. 5Aii, Aiii**). On the other hand, 3 out of 16 PLTX-treated submersion preparations experienced sustained SD without recovery, with reoxygenation unable to restore TPP within the usual 5-minute window (**Table1**, **Fig. 5Ai**).

In animals that experienced post-PLTX azide SD, the general trajectory and the rate of onset appear similar compared to azide control (**Fig. 5Bi, Biv**; SD onset slope, 2 sample Student’s T-test, n = 26, p = 0.9). However, the onset latency of post-PLTX azide SD was much shorter and comparable to PLTX injection SD. (**Fig. 5Biii**; Mann Whitney ranked sum test, n = 26, p < 0.001). Notably, the amplitude of post-PLTX azide SD was significantly reduced by around 36% (**Fig. 5Bv**; Mann Whitney ranked sum test, n = 26, p = 0.005). Corresponding to this observation, PLTX could attenuate and inhibit PLTX injection SD in a dose-dependent manner. Preparations with which azide was unable to induce SD after PLTX treatment had received larger doses of PLTX. (**Fig. 5Bii**; 2 sample Student’s T-test, n = 27, p = 0.006). Similarly, increasing dosage by repeated injections gradually reduced the amplitude of both recoverable and repetitive PLTX-induced SD until the negative DC shift was no longer evident (**Fig. 6Ai, Aii**).

**Figure 6.**
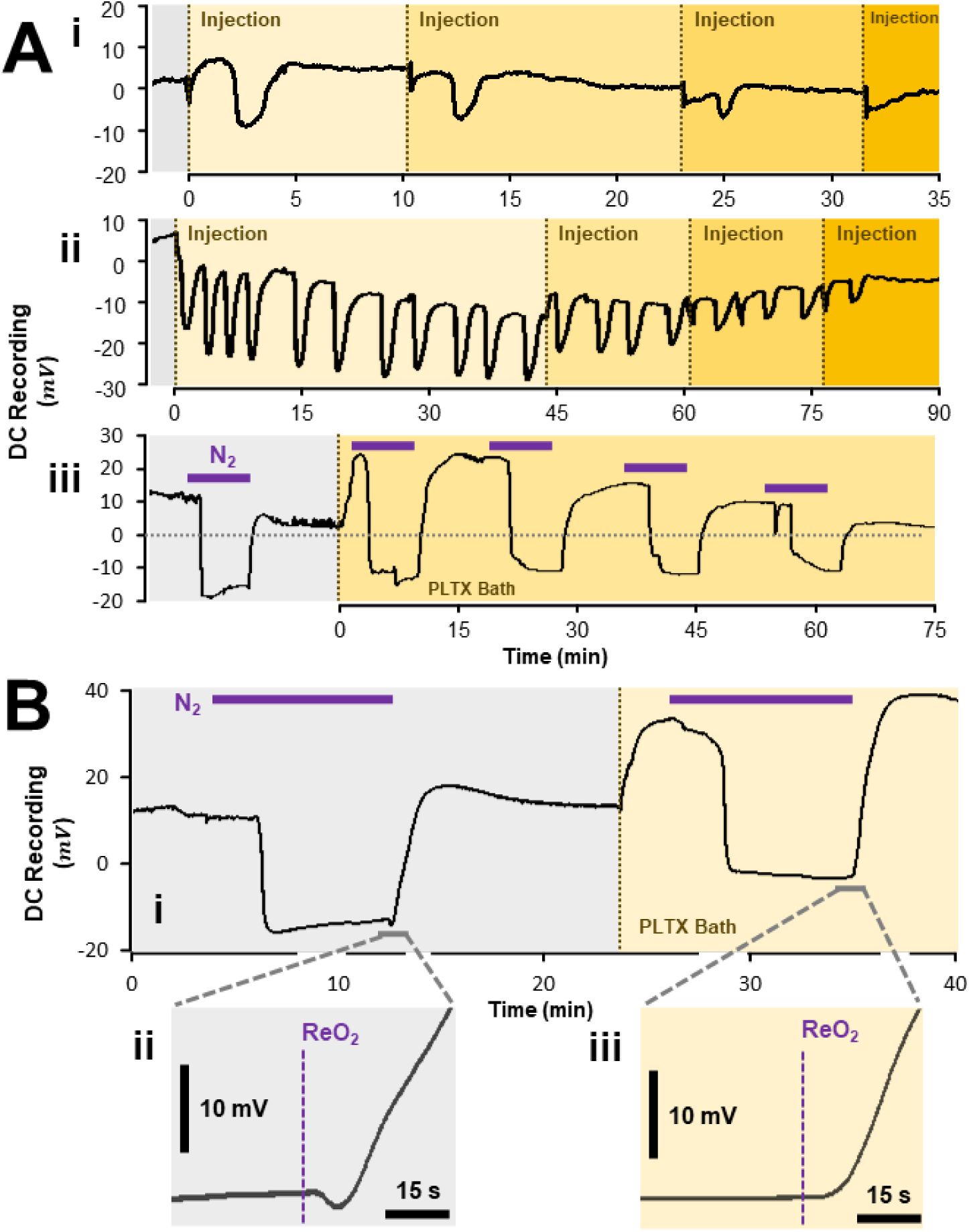
Effects of PLTX on SD attenuation and post-anoxic negativity (PAN). **Ai, ii**: repeated PLTX injections attenuate the amplitude of PLTX-induced SD in a dose-dependent manner. **Aiii**: repeated N_2_ anoxia attenuates anoxic SD amplitude in PLTX bath. **Bi**: comparison of anoxic SD before and after PLTX-bath. **Bii**: PAN occurs immediately following reoxygenation. **Biii:** PLTX bath eradicates PAN. Note the positive shift of TPP following PLTX application. Grey and yellow backgrounds represent untreated and PLTX-treated time intervals, respectively. Purple lines indicate duration of N_2_ anoxia. Vertical dashed lines mark the beginning of treatment. **ReO_2_**: reoxygenation.

Anoxic SD can be initiated in animals where prior PLTX could not initiate SD, suggesting different modifying factors underlying the inhibition of anoxic and PLTX induced SD onset. Additionally, in PLTX bath experiments, it was repeated N_2_ anoxia, rather than increased PLTX concentration, that primarily contributed to the decline in post-PLTX anoxic SD amplitude (**Fig. 6Aiii**).

Finally, it is worth mentioning that PLTX treatment eradicated the post-anoxic negativity (PAN), which is a transient small negative shift of TPP during anoxic SD recovery (**Fig. 6B**). This is an unexpected observation as targeting NKA with ouabain increases the amplitude of the PAN, likely by subtracting the opposing electrogenic effect of the NKA (Van Dusen et al., 2020).

## Discussion

We found that direct injection of PLTX into the locust CNS could initiate SD-like events that are regenerative, all-or-none, sharing similar electrophysiological features with SD evoked by ouabain or azide. PLTX-induced SDs manifests as single, repetitive or sustained forms that appear identical to the various effects of ouabain. In contrast, bath introduction of PLTX failed to generate SD, and there was no indication of an abrupt depression of CNS activity that would signal SD occurring out of the range of our TPP electrode (Leao, 1944; Robertson & Van Dusen, 2021). Given its high molecular weight (2678.5 g/mol, Shimizu, 1983) and the presence of an effective hemolymph-brain barrier in insects (Petschenka et al., 2013), bath-applied PLTX likely could not penetrate into the CNS neuropil and only exerted peripheral effects.

NKA is a fundamental enzyme found in all metazoans (Emery et al., 1998). PLTX exerts its toxic effect by binding to the extracellular surface of the α-subunit of the NKA, at a site that is coupled to but not identical with the binding site for ouabain, and converting the pump into a non-selective cation channel (Rossini and Bigiani, 2011). Vertebrates possess 3 α-subunit isoforms (α1, α2, and α3), with α3 most expressed in the CNS. In contrast, insect pumps have a single isoform that is differentially expressed in the CNS and periphery (Emery et al., 1998; Yang et al., 2019). Despite these differences, insect α-subunits exhibit approximately 80% amino acid sequence similarity with mammals (Emery et al., 1998). Moreover, the pump molecular mechanisms and susceptibility to cardiotonic steroids is conserved across taxa (Blaustein & Hamlyn, 2024). Nevertheless, amino acid substitutions near the ouabain-binding pocket (Dobler et al., 2015; Yang et al., 2019) may account for the reduced sensitivity to PLTX in our experiments compared with mammalian preparations.

Reasonably assuming that the action of PLTX in our experiments was to convert the NKA pump to a channel then such a conversion obviously generates SD. While it remains possible that injected PLTX induces SD through a secondary effect of impaired pump functioning, we propose that, in the present experiments, PLTX likely acts through a pump-to-channel conversion that is well-documented in the literature (DeFelice & Goswami, 2007; Hilgemann, 2003; Rossini & Bigiani, 2011).

### How Does PLTX Generate an SD-like event?

The low latency of PLTX-induced SD aligns with the anticipated effect of pump-to-channel conversion on SD activation. Additionally, the difference in such latency does not seem to be related to the higher potency of PLTX.

Both ouabain and azide trigger SD by inducing pump arrest through direct inhibition or ATP depletion. In this case, the NKA shutdown causes an initially slower accumulation of [Na^+^]_i_ and [K^+^]_o_ until SD initiates. In contrast, PLTX simultaneously arrests pump function and converts it into a large conductance cation channel, by holding open both the Na^+^ and K^+^ gates that normally alternately close during normal pump cycles (Artigas & Gadsby, 2003, 2004). This would immediately activate high Na^+^ influx and K^+^ efflux, thereby reducing SD onset latency. Furthermore, the timing of PLTX-induced SDs was less variable than for ouabain and azide. This difference may arise because intermediate processes between pump arrest and SD ignition are subject to individual variations, such as differing capacities in passive [K^+^]_o_ buffering and potentially, time to hypothetical SD activator release (Andrew, Hartings, et al., 2022; Spong & Robertson, 2013). In contrast, the low variability in the PLTX group suggests that the pathway from induction to activation is brief and comparatively more direct. Indeed, if an endogenous SD activator exists, it should behave similarly to PLTX in this regard.

Accepting that the conversion of NKA to a channel was likely the primary effect of PLTX and main generator of SD, it is worth noting that secondary effects could have modified the nature and severity of the resultant SD. As pumps and channels are structurally and functionally interrelated (Artigas & Gadsby, 2003; DeFelice & Goswami, 2007), the normally tightly coupled transporter gating could be disrupted under pathological conditions in various ways (Clausen et al., 2017; Morth et al., 2008; Ygberg et al., 2021). For example, apart from PLTX, low extracellular pH may also activate a ouabain-sensitive cation conductance in NKA, particularly allowing for the influx of protons and Na^+^ (Vasilyev et al., 2004). Interestingly, such inward current is inhibited under physiological levels of [Na^+^]_o_. Also, different isomeric forms of NKA channels exist in insects and mammals, likely causing different activation and/or conductance properties contributing to SD.

### Inhibition of Anoxic SD

Despite the contrasting success in inducing SD between bath and injected PLTX, both methods disrupted subsequent anoxic SDs. Additionally, in preparations that underwent PLTX-induced SD, a higher dose progressively attenuated the amplitude. These observations suggest that PLTX’s tendency to inhibit anoxic SD occurrence may also interfere with its own ability to sustain SD. In general, the inhibition of hypoxic and anoxic SDs is difficult and often involves potent ion channel inhibitors (Andrew, Farkas, et al., 2022; Hernándéz-Cáceres et al., 1987; Müller, 2000; Müller & Somjen, 1998). However, a key difference exists where unsuccessful inhibition often results in delayed anoxic SD onset in these past studies, yet anoxic SDs are more rapid following PLTX exposure when they are not inhibited. Furthermore, the agents involved in these inhibition studies typically do not initiate SD independently. It is not likely that the lack of SD occurrence is caused by worsening preparation health, as repetitive anoxia does not deteriorate the preparation (Van Dusen et al., 2020) and no apparent perturbations to TPP were observed that would suggest preparation deterioration. Considering SD may be triggered by a PLTX-like endogenous activator (Andrew, Hartings, et al., 2022), and that PLTX occupies a common binding site for pump modulators such as cardiotonic steroids (Artigas & Gadsby, 2004), it is possible that PLTX may compete with the endogenous SD activator for NKA access, or interfere with it functionally. Such interaction could hypothetically reduce the amount NKA involved in channel formation, thus attenuating and preventing SD induction by PLTX itself or by anoxia.

### PLTX Effect on Post-Anoxic Negativity

We found that PLTX eradicated the PAN – a consistent feature of TPP recovery upon reoxygenation, which was originally attributed to activity of the NKA (Spong et al., 2016). However, it is inhibited by bafilomycin (Vacuolar ATPase inhibitor) and not inhibited by ouabain, indicating that it is generated by the electrogenic activity of a V-ATPase (Robertson and Van Dusen, 2021). Indeed, it is enhanced after ouabain treatment due to a reduction of the counteracting electrogenic effect of NKA. The transient negativity of the PAN suggests clearance of protons from the extracellular space. It is larger in males and after more prolonged anoxia. It has been proposed that it reflects proton accumulation due to anaerobic glycolysis during anoxia (Robertson and Van Dusen, 2021). This raises a question of why the PAN is abolished by PLTX and not by ouabain, despite these two agents having the same target.

The most straightforward explanation would be that PLTX also targets the V-ATPase. There have been suggestions that PLTX might convert proton pumps to cation channels (Frelin & Van Renterghem, 1995; Scheiner-Bobis et al., 2002). In addition, PLTX has been associated with effects on development that depend on V-ATPase proton flux (Adams et al., 2006, 2007). In these experiments the PLTX was used to cause cellular depolarization but effects on the V-ATPase cannot be ruled out. Nevertheless, it is generally accepted that the immediate target for PLTX is NKA and any secondary effects are due to ion imbalances (e.g., increased [Na^+^]_i_) or cellular signalling pathways activated by NKA (Rossini & Bigiani, 2011).

### Implications for General SD Mechanisms

Overall, the high degree of similarity between the effects of PLTX and those of ouabain effects suggest that the activation of non-PLTX SD may involve a closely related mechanism. The main argument against NKA as providing the SD-initiating channel is two-fold (Lemale et al., 2022). First, NKA conversion into an open channel is difficult to reconcile with the apparent reversibility of SD, as maintenance of normal pump function is necessary for cellular repolarization and SD recovery. However, we demonstrate that PLTX-induced SD can be reversible *in vivo*. Another point of contention is that the universal tissue distribution of NKA seemingly conflicts with the CNS-restricted occurrence of SD. Yet, the shared susceptibility of insect CNS and rodent brain to PLTX implies that the ubiquity of NKA does not preclude it from exerting CNS-specific effects, such as activating SD. Rather, it is likely that tissue architecture and properties predicate whether or not SD could occur. For instance, the dense aggregation of cellular processes in a volume-restricted space, along with the presence of a diffusion barrier in the CNS may be required for NKA channel to form and drive SD (Robertson et al., 2020).

Critically, the hypothetical SD-initiating channel must close and NKA pump function must be at least partially restored to make SD recovery possible (Lemale et al., 2022). It is still unclear how and why such recovery can occur in the presence of PLTX. One explanation is that PLTX binding to the NKA induces a cation channel reversibly and does not completely abrogate all pump functioning in the neuropil. Detailed *in vitro* characterizations revealed that PLTX could only induce channel formation when NKA is in the E_2_-P conformation (Artigas & Gadsby, 2003, 2004). Conversely, binding under other NKA states does not activate cation conductance nor fully inhibit pump cycling. Furthermore, activated channels do not represent the end-state of the affected pumps, as oscillation between open and closed conformations and PLTX dissociation can still occur (Harmel & Apell, 2006). Alternatively, or perhaps in conjunction, the NKA channel may inactivate to terminate SD. High [K^+^]_o_ during SD may mediate such inactivation, as K^+^ ion were shown to bind and occlude the ion translocation pathway in the pump channel (Artigas & Gadsby, 2003). This occlusion requires a simultaneous decrease in [Na^+^]_o_ and increase in [K^+^]_o_, consistent with the effect of an initial large cationic current during SD. Therefore, the reversal of NKA conformation change, residue pump functions, and channel inactivation potentially allows for SD recovery in the presence of PLTX. The hypothetical SD activator acting through a PLTX-like mechanism may terminate SD in a similar manner.

Despite the marked similarity between the properties of SD in mammals and insects, generalizing the current results should be done cautiously. Between insects and mammals, there are many differences in NKA subunit isoforms, ion homeostatic mechanisms, diffusion barriers, etc. These differences could account for some phenomena we describe that might not be evident in mammalian nervous tissue, like brain slices. For instance, though clustered, repetitive SD is well documented in mammalian brains and ouabain can induce repetitive SD in rat brain (Balestrino et al., 1999), ouabain or PLTX in rodent brain slices usually generates terminal SD, even at low concentrations. Hence, it remains possible that the reversibility and repetitive nature of PLTX-induced SD described here may be peculiar to insect CNS or a function of an intact blood/hemolymph-brain barrier.

## Conclusion

In this study, we present evidence that PLTX can trigger reversible SD in the locust CNS *in vivo*. This SD shares uniform electrophysiological features with those induced by conventional means. Further, we find that PLTX attenuates and prevents the occurrence of anoxic SD, a peculiar phenomenon warranting further investigation. Based on the present observations and well documented biophysical data, we propose that locust SD may be driven by an NKA conversion into a channel in the presence of an endogenous activator *in vivo*.

## Abbreviations

CNS: Central Nervous System
HBB: Hemolymph-Brain Barrier
NKA: Sodium/Potassium ATPase
OUA: Ouabain
PAN: Post Anoxic Negativity
PLTX: Palytoxin
SD: Spreading Depolarization
TP: Transperineurial Potential
V-ATPase: Vacuolar ATPase

## Notes

### Competing Interest Statement

The authors have declared no competing interest.

